# Integrated human-machine interface for closed-loop stimulation using implanted and wearable devices

**DOI:** 10.1101/2022.09.04.506494

**Authors:** Vladimir Sladky, Vaclav Kremen, Kevin McQuown, Filip Mivalt, Benjamin H. Brinkmann, Jamie Van Gompel, Kai J. Miller, Timothy Denison, Gregory A. Worrell

## Abstract

Recent development in implantable devices for electrical brain stimulation includes sensing and embedded computing capabilities that enable adaptive stimulation strategies. Applications include stimulation triggered by pathologic brain activity and endogenous rhythms, such as circadian rhythms. We developed and tested a system that integrates an electrical brain stimulation & sensing implantable device with embedded computing and uses a distributed system with commercial electronics, smartphone and smartwatch for patient annotations, extensive behavioral testing, and adaptive stimulation in subjects in their natural environments. The system enables precise time synchronization of the external components with the brain stimulating device and is coupled with automated analysis of continuous streaming electrophysiology synchronized with patient reports. The system leverages a real-time bi-directional interface between devices and patients with epilepsy living in their natural environment.

## I. Introduction

In recent papers [1], [2], we have described a distributed brain co-processor system coupled with an implantable neurostimulator and sensing device (INSS) that enables long-term wireless streaming of continuous intracranial EEG. The system is linked to the cloud environment that utilizes of high-cost computational algorithms to extract clinically useful information from the collected data. Automated tools for biomarker tracking include interictal epileptiform spike (IES) rates, accurate seizure reports [2], and classification of behavioral states. [3], [4]

A particular challenge with the system is its limitation of data collection using the MS Windows running tablet computer (Epilepsy Patient Assist Device – EPAD) and the clock synchronization between the INSS and EPAD. Here we introduce a system that synchronizes multiple device clocks and enables biomarker tracking that can be used to guide electrical brain stimulation therapy. The integration and clock synchronization of INSS with off-the-body smart commercial electronics (phone and watch) enable analysis of human behavior and biomarkers in behaving humans. The system can be used for clinical and neuroscience research applications, where visual, audio, or motor tasks are presented on a commercial device.[5]–[8] The system consists of two main components. The first one is the investigational Medtronic Summit RC+S™ (RC+S™), a rechargeable sensing and stimulation implantable device with a bi-directional application programming interface. The second component is a smartphone and smartwatch in near proximity to a patient with precise time synchronization of all components using inherent functions of commercial electronics. We validate and test these capabilities in patients with epilepsy living in their home environment.[5]–[7], [9]

## II. Methods

### A. Embedded and Distributed Implantable Brain Stimulation & Sensing System

Four human subjects were implanted with the RC+S™ at Mayo Clinic under an FDA IDE: G180224 and Mayo Clinic IRB: 18-005483 “Human Safety and Feasibility Study of Neurophysiologically Based Brain State Tracking and Modulation in Focal Epilepsy”. The FDA study is https://clinicaltrials.gov/ct2/show/NCT03946618

The patients provided written consent in accordance with the IRB and FDA requirements. The INSS device is an investigational rechargeable system for closed loop stimulation and continuous intracranial electroencephalogram (iEEG) data sensing and telemetry. Patients with implanted INSS device use an antenna and MS Windows running tablet (Fig. 1). The tablet streams information using low energy Bluetooth protocol to a smartphone and smartwatch. The system streams iEEG data, device and system details, patient annotations, including medication, notes, audio/video, and results of surveys up to the cloud for later distributed analytics and clinical and technical team review.

**Figure 1.**
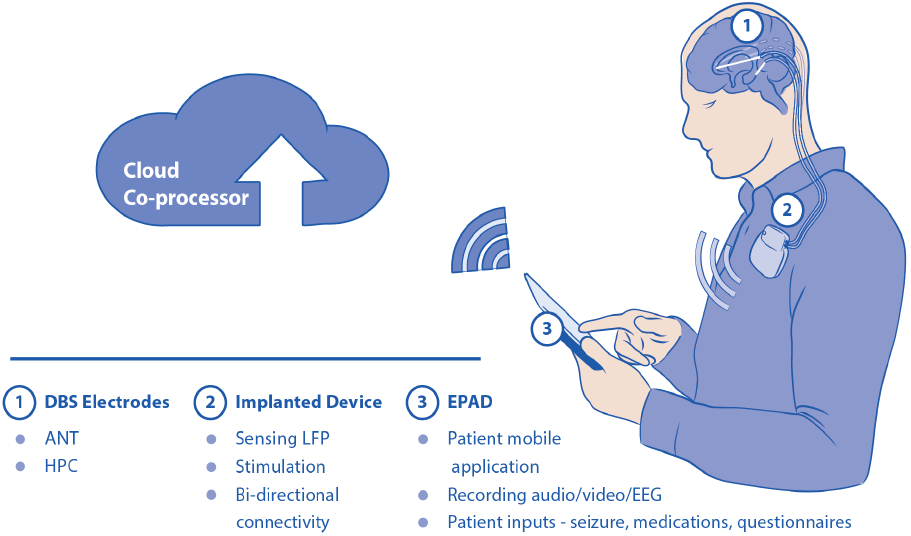
Implantable Neural Sensing & Stimulation System (INSS). DBS electrodes (1) are implanted in the brain (ANT – anterior nucleus of thalamus, HPC – hippocampus) and connected to the rechargeable implanted device (2) enabled for sensing and stimulation and bi-directional connectivity to distributed system (3) Epilepsy Patient Assist Device (EPAD) that is capable of recording patient inputs (seizures, medication, surveys) and audio/video recording. EPAD sends the data to a cloud co-processor for advanced data processing and can close the loop to an implantable system (2) to dynamically adjust parameters of stimulation.

Streams of continuous iEEG data can be analyzed in a cloud environment in a web browser interface. This way, clinical and technical teams remain in the loop using a web-based Epilepsy Dashboard. The Dashboard enables to review biomarker trends (IES rates & seizures), patient annotations (seizures, auras, medication logs), implanted device data (battery status, telemetry, and EBS parameters), and all the data collected using the phone and watch including results of surveys and audio/video files. Backend engine with an integrated machine learning platform for algorithm development and biomarker tracking enables experts to view and perform brisk annotation of events and label the data as well as to view automatically scored behavioral states (e.g., sleep staging). For instance, an expert can confirm that a detected electrophysiological event or patient reported event was a true positive seizure (Fig. 2).

**Figure 2.**
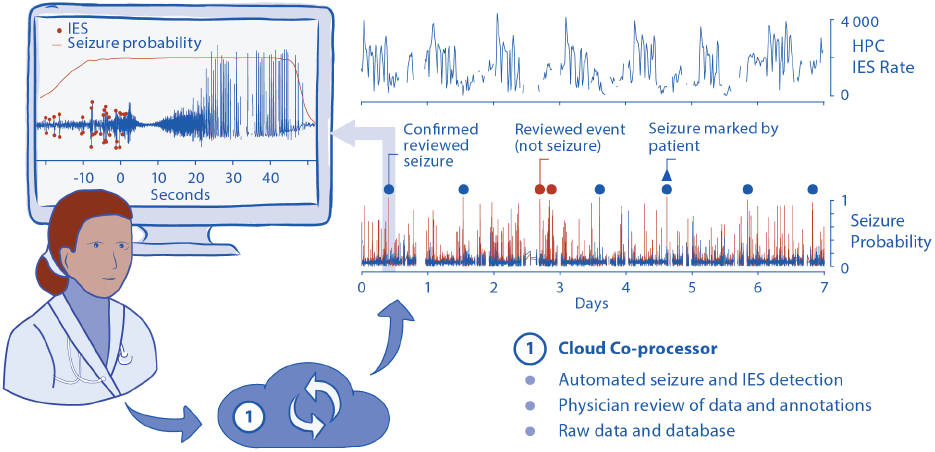
Cloud system coupled with Implantable Neural Sensing & Stimulation System (INSS). Data published to the cloud system can be displayed and reviewed by a physician and technical or clinical team. Automatically detected events (such as seizures or sleep stages) can be assessed and manually corrected to create a ground truth. The advanced data processing in the cloud can enable processing of 24/7 streams of data spanning months and years and create a standard for assessing a patient’s clinical condition. The cloud can then inform the algorithm settings, and if approved by the physician, it can be used to adjust stimulation parameters dynamically. All this enables different stimulation parameters to adapt to changes in brain activity as well as circadian or long-term factors (awake/sleep).

### B. Coupling with Commercial Electronic Devices

The last decade has brought commercial electronics into a state where user experience is smooth and vivid, battery charge lasts long enough, and computational power of the devices enables analytics with high-computational demand to run on the device (smartphone). Moreover, the devices collect vast amounts of physiological data that can be useful for clinical observation or research. Such data includes sensory information (accelerometer, gyroscope, heart rate, blood oxygen saturation, and ECG. Some of the devices utilize embedded algorithms to track and quantify sleep. We are taking advantage of the current commercial technology and synchronizing them with implantable medical devices. This task of synchronization is important to many research applications [10], [11]–[14]. Here we developed a system of commercial electronics (Software tools) that enables us to use it with the implantable RC+S™ system [1], [2], [15]. Fig. 3 shows a physical representation of the system’s components, including implantable benchtop INSS, antenna, 7” MS Windows running tablet, smartphone, and smartwatch.

**Figure 3.**
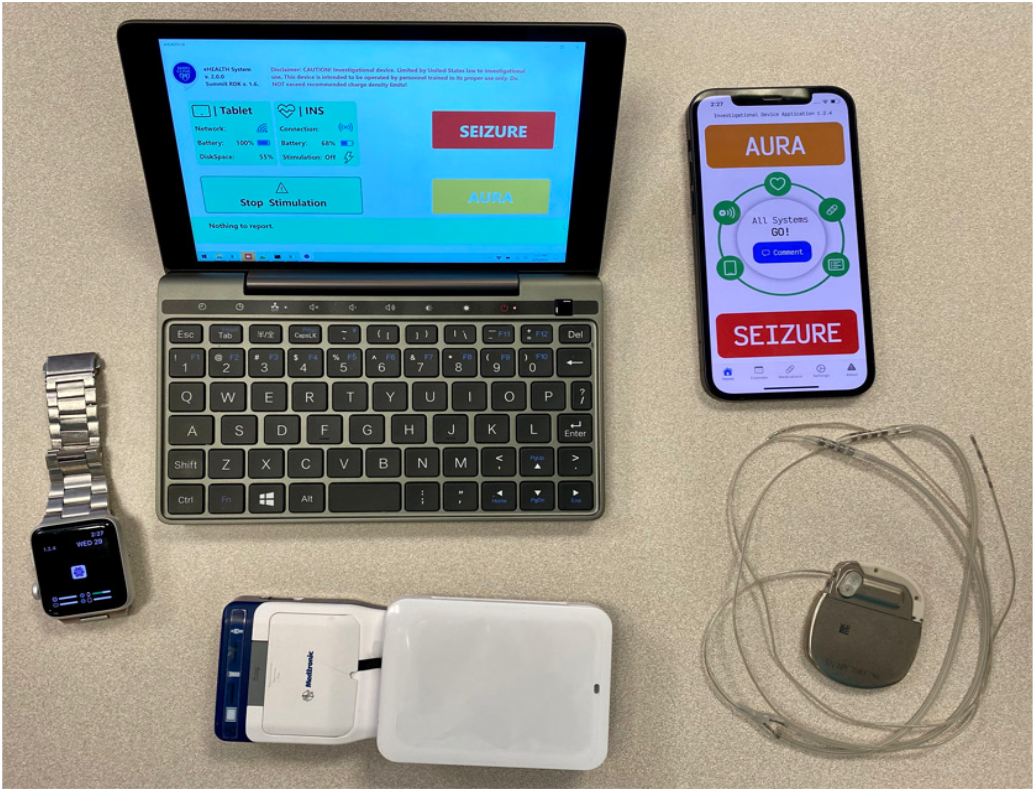
Hardware components of the system, including Implantable Neural Sensing & Stimulation System (INSS), antenna, MS Windows running tablet, smartphone, and smartwatch. A patient has the interface and user experience closest to him on watch, showing the statuses of the devices, including connectivity and batteries. It also enables to launch and ask the patient surveys on watch or phone when the EEG event is detected.

**Figure 4.**
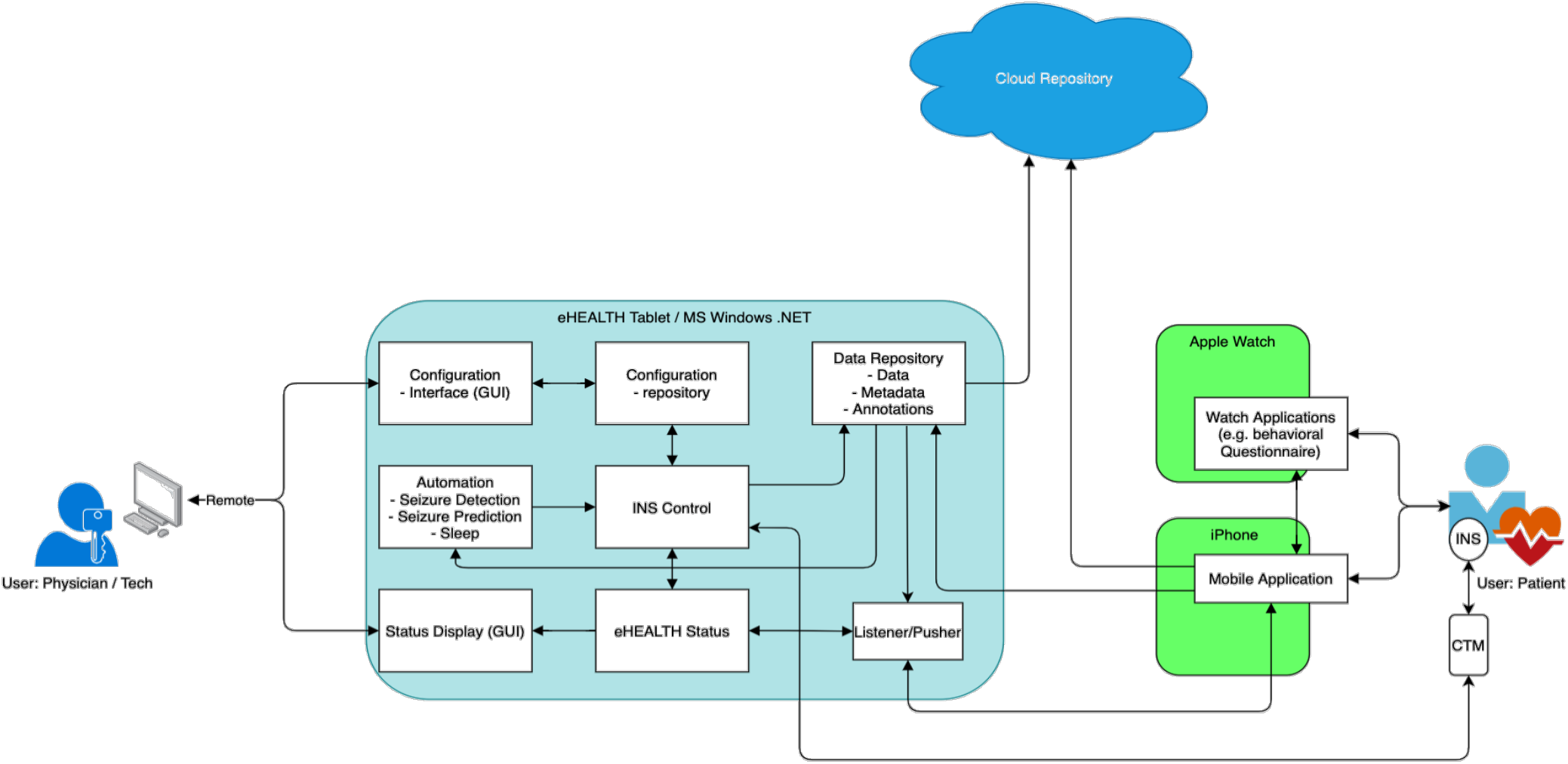
Schematic diagram of the system with essential components and stakeholders (clinical team and patients). A clinical and technical team has a good handle over all technology and a detailed view on settings/data through the whole system, including implantable neurostimulator (INSS) data and control mechanisms. The backbone of MS Windows running tablet computer is the EPAD system connecting via clinician telemetry device (CTM) to the INSS. The original tablet system is now extended to bi-directional connectivity with commercial electronics devices (Apple Watch & iPhone) and includes time synchronization of events and data. The patient experience is brought closer to the patient’s proximity that can perform surveys and tasks, and log all the events (seizures, auras, medication, notes) through the smartphone or smartwatch via a custom-made application. All components are sending anonymized data to the HIPPA compliant cloud system.

Linking the clock synchronization of INSS devices with external devices presents several challenges. To perform experiments in neuroscience (for example, augmented reality task with concurrent iEEG data streaming) with behavioral and external input, the clock synchronization between several device clocks is essential. It should ideally be twice the sampling rate of the fastest signal (in most cases iEEG). We implemented and tested a new method for time synchronization, taking advantage of the INSS feature of monopolar cycling stimulation and the native Apple Watch ECG monitor function.

The INSS device was set in cycling mode (200 msec ON/5 sec OFF) monopolar stimulation with 3 mA, 100 usec pulse width: cathode - anterior nucleus of thalamus, anode – can of the device. We tested a range of stimulation frequencies to evaluate the stimulation artifact visibility using the Apple watch. The INSS device was stimulating and sensing the iEEG as well to be able to synchronize the iEEG data with external sensors/devices. The sampling rate was set to 250 Hz with bandpass filters 0.85 – 100 Hz. We also used an external scalp EEG recording system (Philips EGI High-Density 256 electrode) that was used to capture high density scalp EEG simultaneously to test synchronization capabilities with EEG systems. The sampling rate was set to 1 kHz with 0.5 – 250 Hz bandpass filter and 60 Hz notch filter to remove line noise. In this experiment, we used two more external devices, Apple Watch 7 (iOS 8.5.1) and Apple iPhone 12 (iOS 15.4.1). Apple’s NTP servers synchronize the iPhones and Apple Watches time at “Stratum One” accuracy, within milliseconds of “Stratum Zero” devices.

Timestamps generated by an EndRun Stratum 1 Time Server will typically have 10 microseconds accuracy to UTC and Apple Watch devices might fluctuate about 50 msec around UTC. The Apple Watch team used high-speed cameras to test the accuracy of the Apple Watch. The team filmed the screen at 1,000 frames per second, ensuring that each watch has millisecond precision. The timestamp in msec is given to each ECG record that has been taken on Apple Watch. The ECG record data are stored in HealthKit and linked to our cloud repository in a cloud system (Fig. 1).

This setup ensures that the time stamp of each event on the iPhone and Apple Watch will be within a millisecond precision to the stimulation artifact recorded ECG on Apple Watch in case the artifact is present. As we know, Apple heavily invested in the ECG algorithm, and the data are substantially filtered to record proper ECG. Thus, the essential part of the experiment was to find the best parameters of stimulation to ensure that the stimulation artifact from the INSS monopolar stimulation will be present and visible in the ECG data, so that it can be used later for precise time synchronization of the devices.

## III. Results and discussion

The whole system is now in the field with four patients with temporal lobe epilepsy living in their natural environment. Patients annotate seizure diaries and medication entries and perform mood, memory, and other surveys and tasks. All the data are streamed into the cloud environment and analyzed by automated algorithms [2] and then by manual review of the clinical team. The system battery performance requires charging tablet, phone, and watch at least once per day. The tablet battery (7-10 hours), phone (∼20 hours), and watch (∼24 hours) enable continuous live streaming that is realistically accomplished without severe patient burden. All the components are charged through an external battery pack that can be used in case of backup during longer travels.

Time synchronization between the clock of each individual component is essential for analyzing the data and finding relationships in the multi-modal dataset. We used the inherited Apple Watch application and function to capture ECG, essentially lead II, from one wrist to the other arm by the patient pressing the crown button for 30 sec while Apple Watch is recording the ECG. During this period, the INSS was stimulating to inject the stimulation artifact to the recorded ECG. The example of captured INSS monopolar stimulation in exported pdf file from Apple iPhone is shown in Fig. 5. To be able to synchronize the devices, we tested discrete frequencies at 20, 40, 80, and 125 Hz to find an optimal stimulation frequency that will maximize the visibility of stimulation artifact on Apple ECG to enable better time synchronization. The most prominent stimulation artifact was visible when stimulating at a 40 Hz frequency (Fig. 6). Concurrently, the same stimulation frequency (40 Hz) produced clear detectable artifacts with 200 msec length. We can clearly see the stimulation artifact in Apple Watch ECG, in iEEG, and scalp EEG data (Fig. 6). Considering the fastest sampling frequency 1kHz on scalp EEG data and time synchronization of Apple Watch in msec, we can say that the synchronization can be in the range of 1 msec on scalp EEG and 4 msec on iEEG (250 Hz sampling rate). This can be considered good for performing task experiments and for correlating the data from tasks, surveys, and cameras with iEEG and scalp EEG signals. Such tasks can include verbal memory tasks [16]–[20], navigational memory tasks [10], [12], [13], [21]–[23], including augmented and virtual reality experiments using either augmented reality smart glasses or smartphone/iPad, or surveys and logs taken by patient on the smartphone or smartwatch. This approach provides clock synchronization of EEG recording (including implantable devices) with external devices using commercially available electronics (Apple Watch and iPhone). This approach has the advantage of using external devices to produce an electromagnetic impulse on EEG recording traces for time synchronization [10].

**Figure 5.**
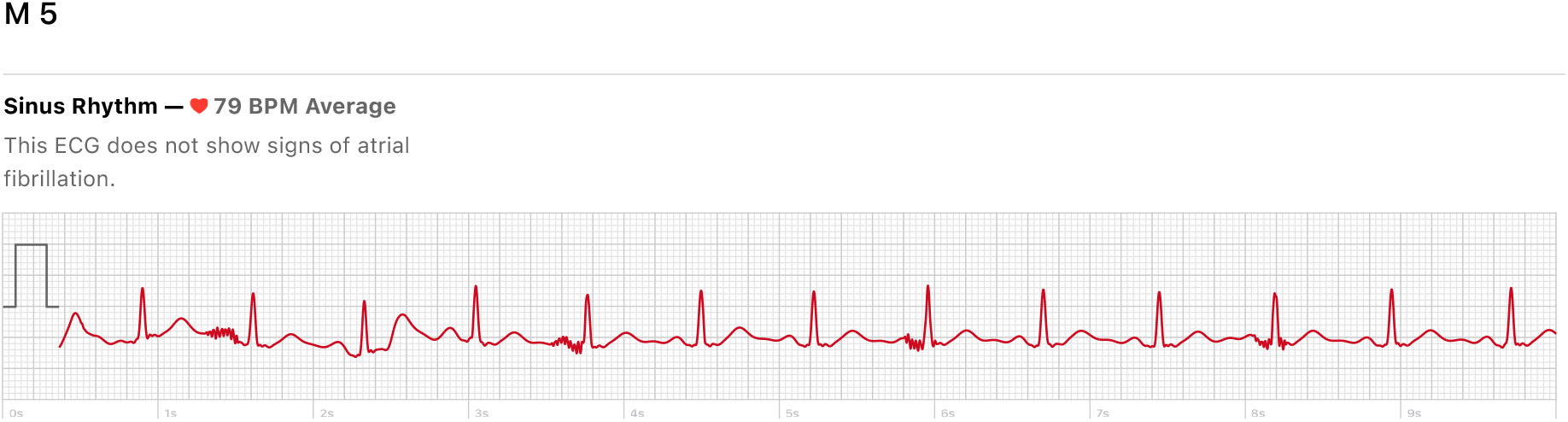
ECG recorded for subject M5 using Apple Watch ECG application as saved in Apple Watch in pdf. The ECG was recorded from the wrist to the other arm (ECG Lead II) while concurrently stimulating by Implantable Neural Stimulator INSS by 40 Hz, 3mA, 100 usec in cycling mode 200 usec ON / 5 sec OFF. Here 10 sec of 30 second native Apple Watch ECG is shown.

**Figure 6.**
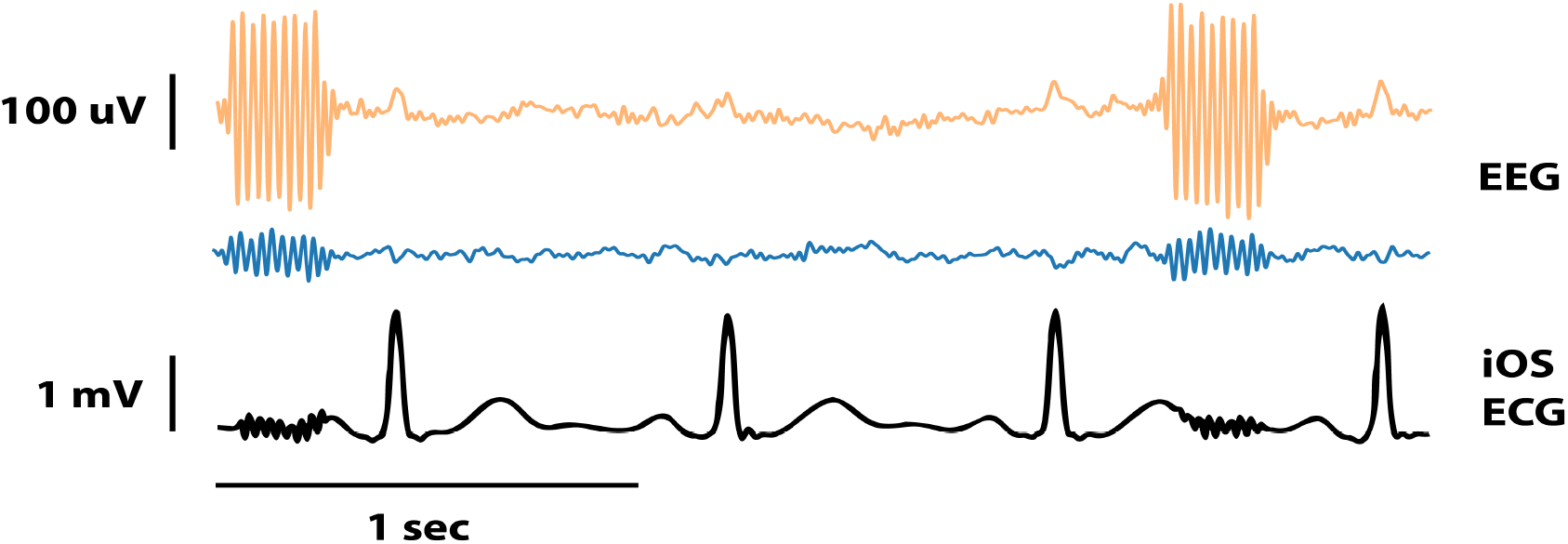
Time synchronization of Implantable Neural Sensing & Stimulation device (INSS) with commercial electronics (smartphone and smartwatch) and external scalp EEG system. The stimulator is cycling high frequency stimulation in the monopolar regime. This can be precisely (in msec) recorded on Apple Watch ECG (black bottom), Internal INSS iEEG (blue middle) and external Philips EGI EEG scalp recording system (top yellow). Stimulation artifacts created by the INSS monopolar stimulation 40 Hz then enable synchronizing clock on all external devices including commercial electronics – watch & phone.

## IV. Conclusions

For the very first time, we show the human-machine system that includes implantable neural sensing & stimulation device coupled with commercial electronics (Apple Watch & iPhone) synchronized in milliseconds precision with intracranial EEG recorded by implanted device as well as external EEG devices measuring concurrently scalp EEG. Such a system can be used for chronic neural sensing and dense behavioral tracking to optimize adaptive stimulation of the brain. Due to highly synchronized clock precision, behavioral testing can include memory tasks, augmented reality tasks, and surveys triggered by events detected in EEG.

## Acknowledgment

We would like to thank all colleagues and the team that created the initial version of EPAD system [1], [15], in particular the programmers Tal Pal Attia, Dan Crepeau and also study coordinator Karla Crocket and EEG technician Cindy Nelson who diligently and relentlessly take care of our patients.

## References

[1] V. Kremen, B. H. Brinkmann, I. Kim, H. Guragain, M. Nasseri, A. L. Magee, T. Pal Attia, P. Nejedly, V. Sladky, N. Nelson, S.-Y. Chang, J. A. Herron, T. Adamski, S. Baldassano, J. Cimbalnik, V. Vasoli, E. Fehrmann, T. Chouinard, E. E. Patterson, B. Litt, M. Stead, J. Van Gompel, B. K. Sturges, H. J. Jo, C. M. Crowe, T. Denison, and G. A. Worrell, “Integrating Brain Implants With Local and Distributed Computing Devices: A Next Generation Epilepsy Management System,” IEEE J. Transl. Eng. Heal. Med., vol. 6, pp. 1–12, 2018.

[2] V. Sladky, P. Nejedly, F. Mivalt, B. H. Brinkmann, I. Kim, E. K. St. Louis, N. M. Gregg, B. N. Lundstrom, C. M. Crowe, T. P. Attia, D. Crepeau, I. Balzekas, V. S. Marks, L. P. Wheeler, J. Cimbalnik, M. Cook, R. Janca, B. K. Sturges, K. Leyde, K. J. Miller, J. J. Van Gompel, T. Denison, G. A. Worrell, and V. Kremen, “Distributed brain co-processor for tracking spikes, seizures and behavior during electrical brain stimulation,” Brain Commun., May 2022.

[3] F. Mivalt, V. Kremen, V. Sladky, I. Balzekas, P. Nejedly, N. M. Gregg, B. N. Lundstrom, K. Lepkova, T. Pridalova, B. H. Brinkmann, P. Jurak, J. J. Van Gompel, K. Miller, T. Denison, E. K. St. Louis, and G. A. Worrell, “Electrical brain stimulation and continuous behavioral state tracking in ambulatory humans,” J. Neural Eng., vol. 19, no. 1, 2022.

[4] V. Kremen, B. H. Brinkmann, J. J. Van Gompel, S. (Matt) M. Stead, E. K. St Louis, and G. A. Worrell, “Automated Unsupervised Behavioral State Classification using Intracranial Electrophysiology,” J. Neural Eng., Oct. 2018.

[5] S. Stanslaski, J. Herron, E. Fehrmann, R. Corey, H. Orser, E. Opri, Kremen, B. Brinkmann, A. Gunduz, K. Foote, G. Worrell, and T. Denison, “Creating neural ‘co-processors’ to explore treatments for neurological disorders,” in 2018 IEEE International Solid - State Circuits Conference - (ISSCC), 2018, pp. 460–462.

[6] V. Kremen, B. H. Brinkmann, I. Kim, H. Guragain, M. Nasseri, A. L. Magee, T. Pal Attia, P. Nejedly, V. Sladky, N. Nelson, S.-Y. Chang, J. A. Herron, T. Adamski, S. Baldassano, J. Cimbalnik, V. Vasoli, E. Fehrmann, T. Chouinard, E. E. Patterson, B. Litt, M. Stead, J. Van Gompel, B. K. Sturges, H. J. Jo, C. M. Crowe, T. Denison, and G. A. Worrell, “Integrating Brain Implants With Local and Distributed Computing Devices: A Next Generation Epilepsy Management System,” IEEE J. Transl. Eng. Heal. Med., vol. 6, pp. 1–12, 2018.

[7] R. Gilron, S. Little, R. Perrone, R. Wilt, C. De Hemptinne, S. Maria, C. A. Racine, S. Wang, J. L. Ostrem, P. S. Larson, D. Doris, N. B. Galifianakis, I. Bledsoe, M. S. Luciano, H. E. Dawes, A. Gregory, V. Kremen, D. Borton, T. Denison, and P. A. Starr, “Chronic wireless streaming of invasive neural recordings at home for circuit discovery and adaptive stimulation,” bioarxiv, 2020.

[8] T. Pal Attia, D. Crepeau, V. Kremen, M. Nasseri, H. Guragain, S. W. Steele, V. Sladky, P. Nejedly, F. Mivalt, J. Herron, M. Stead, T. Denison, G. A. Worrell, and B. H. Brinkmann, “Epilepsy Personal Assistant Device -A Mobile Platform for Brain State, Dense Behavioral and Physiology Tracking and Controlling Adaptive Stimulation,” Front. Neurol., 2021.

[9] D. A. Borton, H. E. Dawes, G. A. Worrell, P. A. Starr, and T. J. Denison, “Developing Collaborative Platforms to Advance Neurotechnology and Its Translation,” Neuron. 2020.

[10] M. Stangl, U. Topalovic, C. S. Inman, S. Hiller, D. Villaroman, Z. M. Aghajan, L. Christov-Moore, N. R. Hasulak, V. R. Rao, C. H. Halpern, D. Eliashiv, I. Fried, and N. Suthana, “Boundary-anchored neural mechanisms of location-encoding for self and others,” Nat. 2020 5897842, vol. 589, no. 7842, pp. 420–425, Dec. 2020.

[11] J. Miller, A. J. Watrous, M. Tsitsiklis, S. A. Lee, S. A. Sheth, C. Schevon, E. H. Smith, M. R. Sperling, A. Sharan, A. A. Asadi-Pooya, G. A. Worrell, S. Meisenhelter, C. S. Inman, K. A. Davis, Lega, P. A. Wanda, S. R. Das, J. M. Stein, R. Gorniak, and J. Jacobs, “Lateralized hippocampal oscillations underlie distinct aspects of human spatial memory and navigation,” Nat. Commun., vol. 9, no. 1, 2018.

[12] J. Park, H. Lee, T. Kim, G. Y. Park, E. M. Lee, S. Baek, J. Ku, I. Y. Kim, S. I. Kim, D. P. Jang, and J. K. Kang, “Role of low- and high-frequency oscillations in the human hippocampus for encoding environmental novelty during a spatial navigation task,” Hippocampus, vol. 24, no. 11, pp. 1341–1352, Nov. 2014.

[13] H. Eichenbaum, “The role of the hippocampus in navigation is memory,” J. Neurophysiol., vol. 117, no. 4, pp. 1785–1796, Apr. 2017.

[14] S. Maidenbaum, A. Patel, I. Garlin, and J. Jacobs, “Studying Spatial Memory in Augmented and Virtual reality,” bioarxiv, 2019.

[15] T. P. Attia, D. Crepeau, V. Kremen, M. Nasseri, H. Guragain, M. Stead, T. Denison, G. A. Worrell, and B. H. Brinkmann, “Epilepsy Personal Assistant Device — A Mobile Platform for Brain State, Dense Behavioral and Physiology Tracking and Controlling Adaptive Stimulation,” Front. Neurosci., vol. 12, no. July, pp. 1– 14, 2021.

[16] M. T. Kucewicz, B. M. Berry, L. R. Miller, F. Khadjevand, Y. Ezzyat, J. M. Stein, V. Kremen, B. H. Brinkmann, P. Wanda, M. R. Sperling, R. Gorniak, K. A. Davis, B. C. Jobst, R. E. Gross, B. Lega, J. Van Gompel, S. M. Stead, D. S. Rizzuto, M. J. Kahana, and G. A. Worrell, “Evidence for verbal memory enhancement with electrical brain stimulation in the lateral temporal cortex,” Brain, Jan. 2018.

[17] K. V Saboo, I. Balzekas, V. Kremen, Y. Varatharajah, M. Kucewicz, R. K. Iyer, and G. A. Worrell, “Leveraging electrophysiologic correlates of word encoding to map seizure onset zone in focal epilepsy : Task- - dependent changes in epileptiform activity, spectral features, and functional connectivity,” Epilepsia, vol. 00, pp. 1–13, 2021.

[18] V. S. Marks, K. V Saboo, M. Lech, T. P. Thayib, P. Nejedly, V. Kremen, G. A. Worrell, and M. T. Kucewicz, “NeuroImage Independent dynamics of low, intermediate, and high frequency spectral intracranial EEG activities during human memory formation,” vol. 245, no. October, 2021.

[19] M. T. Kucewicz, J. Dolezal, V. Kremen, B. M. Berry, L. R. Miller, A. L. Magee, V. Fabian, and G. A. Worrell, “Pupil size reflects successful encoding and recall of memory in humans,” Sci. Rep., vol. 8, no. 1, 2018.

[20] M. T. Kucewicz, J. Cimbalnik, J. Y. Matsumoto, B. H. Brinkmann, M. R. Bower, V. Vasoli, V. Sulc, F. Meyer, W. R. Marsh, S. M. Stead, and G. A. Worrell, “High frequency oscillations are associated with cognitive processing in human recognition memory,” Brain, vol. 137, no. 8, pp. 2231–2244, 2014.

[21] E. K. Miller, M. Lundqvist, and A. M. Bastos, “Working Memory 2.0,” Neuron, vol. 100, no. 2, pp. 463–475, 2018.

[22] J. Jacobs, “Hippocampal theta oscillations are slower in humans than in rodents: Implications for models of spatial navigation and memory,” Philos. Trans. R. Soc. B Biol. Sci., vol. 369, no. 1635, Feb. 2014.

[23] Y. Pu, B. R. Cornwell, D. Cheyne, and B. W. Johnson, “The functional role of human right hippocampal/parahippocampal theta rhythm in environmental encoding during virtual spatial navigation,” Hum. Brain Mapp., vol. 38, no. 3, pp. 1347–1361, Mar. 2017.

